# A Deep Learning Model for Quantifying and Mapping Human Spiral Ganglion Neurons (SGNs): The SGN Base Model

**DOI:** 10.64898/2026.11.10.765481

**Authors:** Abbie K. Hall, Haobing Wang, Nicole Kim, Christopher K. Giardina, Julie G. Arenberg, Alicia M. Quesnel

**Author notes:** Corresponding Author: Alicia M. Quesnel, MD.

## Abstract

**Introduction:** Spiral ganglion neurons (SGNs) transmit auditory information and are a histological marker that have been quantified for decades to study various hearing loss pathologies in human temporal bone (hTB) specimens. Historically, quantification of SGNs has been labor-intensive and subject to manual bias and error. With the advent of digital microscopy, this study introduces and assesses the accuracy of a deep learning model for SGN quantification, providing an efficient approach that enables in-depth analysis of SGN distribution through three-dimensional frequency mapping.

**Methods:** Digitized mid-modiolar slides from six hTBs from the Mass Eye and Ear Otopathology Laboratory Collection, representing a range of ages, post-mortem times, and diagnoses, were selected to train a computer vision model. Using Dragonfly 2024.1, the nucleus and cytoplasm of each SGN were manually segmented against background and used as the training corpus. The model was systemically evaluated in eighteen new cases to assess its accuracy across diverse histopathological presentations. For the specimen with the lowest performance, five small regions were manually annotated to augment the training dataset to evaluate the adaptability of the model. Next, the model was applied to every available slide that includes Rosenthal’s canal for a single case to assess model performance throughout the entire cochlea. Model performance was evaluated by comparing model SGN counts with counts by an experienced investigator, classifying each model prediction as a true positive, false positive, or false negative. Using coordinates output by the model, three-dimensional reconstructions were generated to analyze model performance from base to apex.

**Results:** A U-Net convolutional neural network was developed with an average Dice coefficient of 0.91. Across eighteen cases representing diverse SGN morphologies and densities, the model achieved an F1-score of 0.91. For the case with the lowest performance, fine-tuning the model by adding additional training data from the same case improved the r^2^ value from 0.919 to 0.996. When applied to all available slides from a single case, the model achieved an F1-score of 0.96 across the entire cochlea.

**Conclusion:** We present the pretrained SGN Base Model: an approach for quantifying SGNs in hTB specimens that improves efficiency while maintaining high accuracy in comparison to manual quantification. The model performs robustly throughout the cochlea and is adaptable to a diverse range of archival hTB specimens with variable SGN densities and morphologies. Model-generated spatial coordinates for each SGN enable three-dimensional analysis and detailed mapping of SGN density from base to apex.

## Introduction

Spiral ganglion neurons (SGNs) are necessary for sound processing and perception, transmitting auditory information from the inner ear to the central nervous system. Degeneration of SGNs has been associated with hearing loss and is a pathological feature observed in aging, noise exposure, ototoxicity, and genetic hearing loss (1–5). Additionally, these auditory neurons are the primary targets for hearing restoration via electrical stimulation from cochlear implants, bypassing mechanical hair cell transducers which rely on functional acoustic to electric stimulation to generate an acoustic precept. As the primary electric relay, SGNs have been studied and quantified to better understand the mechanisms of a broad range of diagnoses and assist in outcomes of cochlear implantation (1,6–8). Although animal models have been useful in characterizing function and degeneration of SGNs, there are interspecies differences in quantity and three-dimensional distributions of SGNs relative to humans (9–13). Additionally, SGNs degenerate at much faster rates in animal models in comparison to humans (14).

Human studies of SGN quantification are based solely on post-mortem human temporal bone (hTB) specimens due to the inability to quantify SGNs by any method other than tissue analysis. Historically, SGNs in hTBs have been manually quantified using hematoxylin and eosin (H&E) staining under light microscopy, a very tedious and time-consuming process that is prone to inter-rater variance given that normal-hearing individuals may have, on average, more than 30,000 neurons (15–17). Stereology was introduced to improve efficiency; however, while this approach accurately estimates the total number of SGNs, it is unable to analyze the distribution throughout the entire cochlea and does not preserve the spatial arrangement of SGNs throughout the cochlear spiral (18,19).

With the advent of digital microscopy, scans and automated pipelines (i.e.: computer vision and machine learning) have been widely applied throughout molecular biology research to identify a variety of cell types (20–24). In addition to the speed of quantification vs hand-counting, automated methods also help to standardize analysis by reducing inter-rater variance. More recent computer vision approaches utilize deep learning models (UNet, UNet++) and have been previously used to quantify SGNs in human and animal specimens, ultimately improving efficiency while maintaining high accuracy (25–27). Although the previously published machine learning model for SGN quantification in hTBs developed at Mass Eye and Ear demonstrated strong performance, its generalizability across specimens may be limited by the morphological heterogeneity of SGNs. Neuronal density varies among individuals based on etiology and age (1,2,28–30). Additionally, the appearance of these neurons, including color, size, and shape, may differ due to processing techniques, staining intensity, and post-mortem cellular changes (31). The previously published model was trained and validated using specimens with standard age-related hearing loss while this present study expands upon this framework by incorporating automated SGN identification across a broader range of pathologies, post-mortem intervals, and storage times. By incorporating segmentations of both SGN nuclei and cytoplasm, the model described here represents a preliminary step toward developing a more generalizable SGN quantification approach that is less dependent on specific etiologies of hearing loss or post-mortem intervals. Furthermore, the model was developed using freely available software, facilitating accessibility and adaptability for further applications.

Here, we introduce and describe an adaptable deep learning model that efficiently and accurately quantifies SGNs in a variety of hTBs with different staining intensities, post-mortem times, and neuronal densities. The SGN Base Model has been rigorously compared to gold-standard methods for quantification and has the potential to be applied to thousands of archival hTBs ready for analysis in the Mass Eye and Ear hTB collection and outside collections.

## Methods

### Specimen Processing and Selection

The specimens used in this study are from the Mass Eye and Ear Otopathology Laboratory Collection. Each hTB was processed using standard histological techniques, consisting of fixation, decalcification, celloidin-embedment, and sectioning to ultimately produce about 400 horizontally sectioned 20 µm thick slices (32). Every tenth section was stained using H&E, producing about 40 stained slides for analysis.

hTBs selected to train the model represented a range of diagnostic categories: normal, sensorineural hearing loss, genetic hearing loss, otosclerosis, Meniere’s disease, and post-cochlear implantation (Table 1). These specimens consisted of variable post-mortem times, ages, and years of processing (year of death) to reflect the heterogeneity of SGN morphology and density, with the purpose of testing the model’s applicability to a wide variety of cases. All protocols for this study were approved by the Institutional Review Board of Mass Eye and Ear (IRB #2021P001593).

**Table 1.** Cases selected to train the model considering different variables that would affect SGN density and morphology, including diagnosis, age, post-mortem time, and storage time.

| **Case** | **Diagnosis** | **Age**<br>**(Years)** | **Post-Mortem Time**<br>**(Hours)** | **Year of Death** |
|---|---|---|---|---|
| 1 | Meniere’s Disease | 63 | 5 | 1969 |
| 2 | Otosclerosis | 72 | 23 | 1980 |
| 3 | Sensioneural Hearing Loss | 92 | 9.5 | 2010 |
| 4 | Genetic (X-linked) | 80 | 19.5 | 2015 |
| 5 | Normal | 24 | 7.5 | 2017 |
| 6 | Cochlear Implant | 90 | 53 | 2020 |

### Slide Digitization and Cropping

Each archival slide was scanned using a Leica Aperio AT2 scanner (Leica Microsystems, Wetzlar, Germany) with a 20x objective (NA 0.75) to generate digitized slides, or “eSlides.” The scanner acquired standard pyramid titled SVS files using the best focal plane to produce high resolution images of 0.501 µm or 0.503 µm per pixel.

Following digitization, eSlides were analyzed and cropped within Aperio ImageScope (Leica Microsystems, Wetzlar, Germany). A mid-modiolar slide from each case selected to train the model was manually cropped around the cochlea, generating TIFFs in uniform dimensions of 20,0000 pixels x 10,000 pixels, allowing them to be imported as a single image stack (Fig. 1a).

**Fig. 1.**
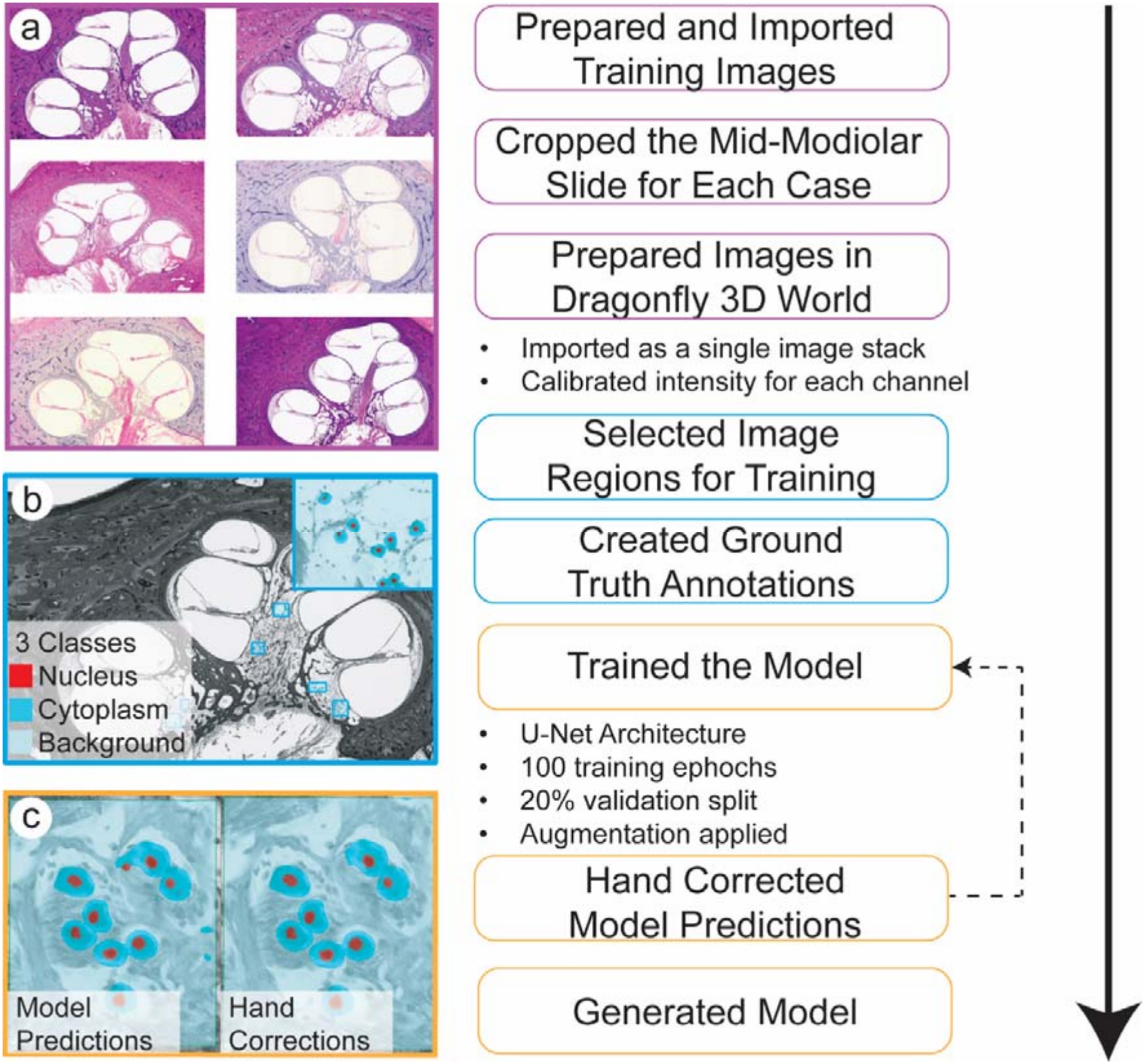
Building the model. Cropped high-resolution mid-modiolar slides for the six cases selected to train the model (a). The segmented areas for one of the training cases, with a magnified view an area containing all three classes: nucleus (red), cytoplasm (cyan), and background (light blue) (b). Initial model predictions (left) and the corresponding hand-corrected predictions (right) used for subsequent rounds of training.

### Training Set Annotation

The six cropped images of equal dimensions were imported into Dragonfly 3D World 2024.1 (Comet Software, Montreal, QC, CA) as a single image stack using all three-color channels: red, green, and blue. Each channel was calibrated using a custom-defined dimensional unit, ranging from 0 to 100, with 0 corresponding to the lower peak and 100 to the higher peak of the intensity histogram.

Small image frames from each slide were hand-annotated using Segmentation Wizard in Dragonfly 3D World (Fig. 1b). These frames were created in various cochlear regions to capture different neuronal densities and cochlear turns, ensuring diversity in the training set. Pixels within the hand-annotated frames were assigned to one of three classes: nucleus, cytoplasm, and background. To improve model discrimination, areas containing bone, Schwann cells, vessels, and artifacts were partially included within the annotated frames and annotated as background.

A U-Net convolutional neural network was adapted from the built-in deep learning framework for semantic segmentation in Dragonfly 3D World and was implemented for three-class semantic segmentation. The network consisted of five encoding-decoding levels with an initial filter count of 64, doubling down at each down sampling step. Training was performed on two-dimensional (2D) image patches of 256 x 256 pixels and all three-color channels; red, green, and blue, were used as data input. For each cycle, the model was trained for 100 epochs with a batch size of 64. Categorical cross-entropy loss was optimized using the Adadelta optimizer with a learning rate of 1.0. Data augmentation was applied tenfold, including horizontal and vertical flips, rotations, zooming, and shear transformations.

The model was trained using the annotated frames as input. Following an initial round of training, the model generated predications in newly selected frames. These predictions were hand-corrected and incorporated as additional training data (Fig. 1c). This process was repeated, resulting in 5-6 annotated frames per case. Using this approach, the model iteratively learned from incorrect predictions and ultimately improved its accuracy and robustness across different cochlear regions and neuronal densities.

### Applying the Model

Each digitized slide selected to validate the model was cropped and saved as a high-resolution TIFF with equal dimensions (Fig. 2a). After importing into Dragonfly 3D World as a single image stack, each color channel was calibrated, and Rosenthal’s canal was segmented to create a binary mask to define a region of interest (ROI) to apply the model (Fig. 2b-d). The trained deep learning model was applied within this mask to classify each voxel into one of three classes: nucleus, cytoplasm, or background (Fig. 2e,f). For the lowest performing case, the model was fine-tuned using a subset of manually annotated areas from the same specimen, followed by reapplication and reassessment, to determine the adaptability of the model (Fig. 2g). Voxels assigned to the nucleus class were exported as a discrete ROI and further analyzed using the Connected Components algorithm in Dragonfly 3D World, which separated individual nuclei and enabled quantitative measurements, including surface area and spatial coordinates (Fig. 2h). Using surface area, a size-cut off was applied to exclude false positives and false negatives caused by background noise (Fig. 2i). The output of the model can be hand-corrected to add missed SGNs, and remove mislabeled structures, such as bone, blood vessels, Schwann cells, and artifacts (Fig. 2j). However, for the purpose of validating the model, no hand-correction was performed.

**Fig. 2.**
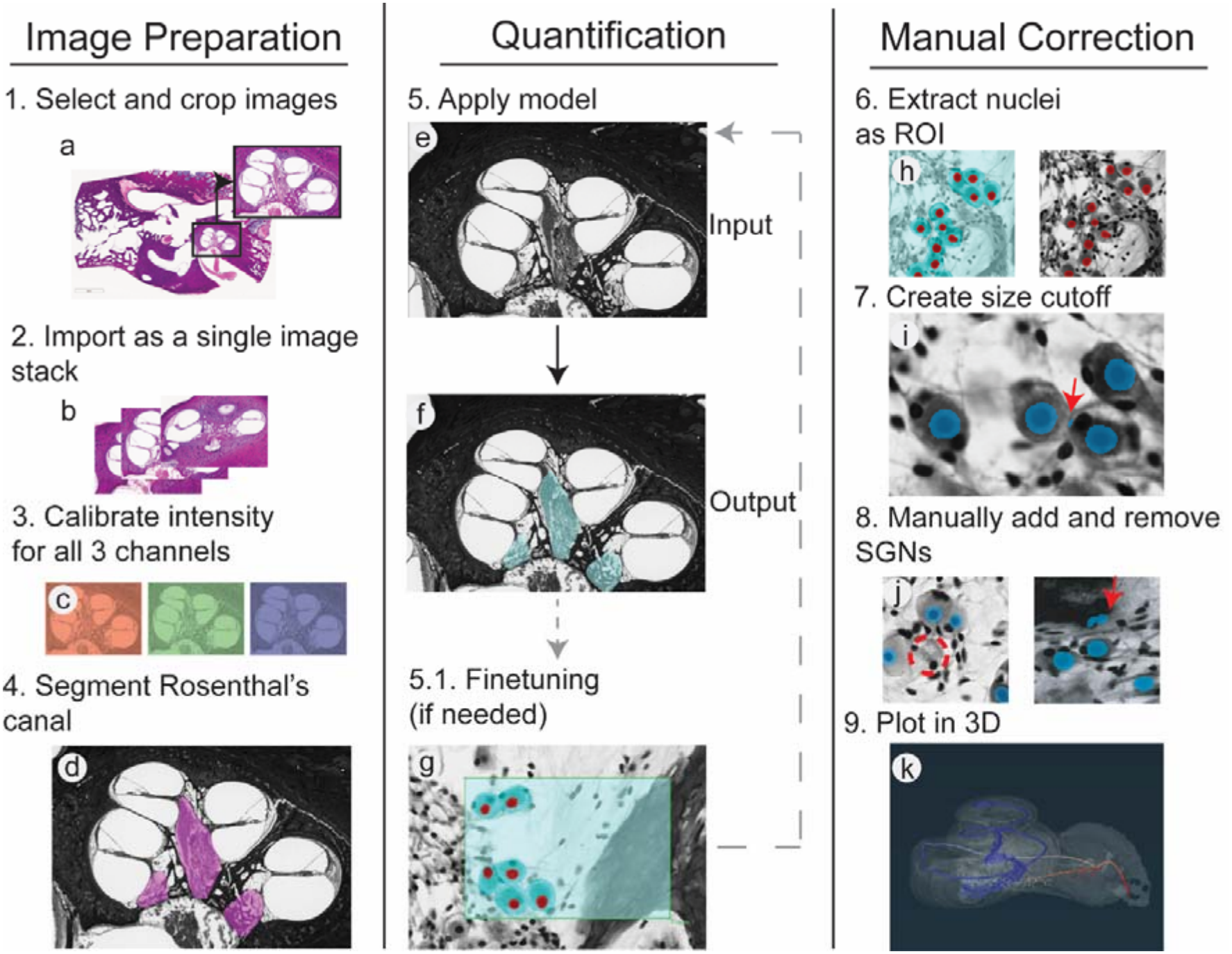
Applying the model. Each digitized slide cropped (a) and imported (b) into Dragonfly 3D World. The image intensity calibrated for all three channels: red, green, and blue (c). Rosenthal’s canal segmented, creating a mask to apply the model (d). The input (e) and output (f) of the model. A small area used as additional training data to fine-tune the model (g). A magnified view of the three classes: nucleus (red), cytoplasm (cyan), and background (red) (h). Background noise (red arrow) eliminated by using a size cutoff (i). A missed SGN (dashed red circle), and bone mislabeled as an SGN (red arrow) (j). SGN coordinates output from the model plotted in 3D (k).

### Mapping SGNs

First, all eSlides containing the cochlear region were cropped and down sampled in the X and Y dimensions to a resolution of 2.5 µm x 2.5 µm per pixel, while maintaining the original Z spacing of 200 µm. The cropped and down-sampled images were then imported into Amira3D 2024.1 (Thermo Fisher Scientific, Waltham, MA, USA) and aligned using the “AlignSlices” function. The X, Y, and Z coordinates of the SGNs were exported as a csv file from Dragonfly 3D World and imported into Amira3D. The SGN coordinates were aligned according to the 2D affine transformation generated by “AlignSlices,” using a custom Amira Python script.

After aligning the slices and SGNs, the spiral path along Rosenthal’s canal was constructed from the cochlear base along with the canal’s centerline toward the apex. Similarly, the spiral path of the organ of Corti was constructed by marking the intersection points of the inner and outer pillar cells also from base to apex. Both spiral pathways were created in Amira3D as B-spline curves and uniformly sampled at 600 points. For each point along the Rosenthal’s canal and organ of Corti pathways, an associated characteristic Frequency (CF) was calculated based on the formulas derived from 3D synchrotron imaging (33). Each SGN was assigned a CF by finding its nearest point (smallest Euclidian distance) on Rosenthal’s canal pathway (Fig. 2k).

Equation 1: Assigning CF by Percent Distance along Organ of Corti

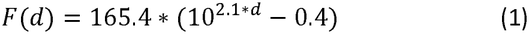

Equation 2: Assigning CF by Percent Distance along Rosenthal’s Canal

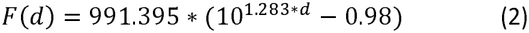

### Determining Modiolar Axis and Assigning SGN Angular Insertion Depth

The modiolar axis was identified on the mid-modiolar section near the most apical point of Rosenthal’s canal and designated as the centerline of the modiolus. The axis served as the rotation axis for quantifying rotation angles of points along the Rosenthal’s canal and organ of Corti pathways, as well as the SGNs. The spatial coordinates of these points were projected onto a two-dimensional plane orthogonal to the rotation axis, and the rotation angles were determined on this projection plane, with the position of the round window membrane defined as 0°. Similar to assigning a CF to each SGN based on the CF of the nearest Rosenthal’s canal segment, each SGN was assigned a modiolar angle corresponding the angle of the closest Rosenthal’s canal segment. The modiolar angle calculations were performed using custom Python scripts. Histograms of SGN count as function of modiolar angle and percent distance along Rosenthal’s canal were generated using Python.

### Assessment of Model Performance

Model performance was evaluated by comparing automated SGN counts by the generic model to manual SGN counts by an experienced investigator. SGNs were manually counted using Fiji (ImageJ) (34). Coordinates for each manually counted SGN were imported into Dragonfly 3D World and overlaid with the model’s predictions. Each predicted SGN by the model was classified as a true positive (TP), false positive (FP), or false negative (FN) relative to manual SGN counts. Model performance metrics, such as precision, recall, and F1 score, were calculated to assess accuracy. Additionally, a regression analysis was conducted to assess concordance between model counts and manual counts.

Equation 3: Recall

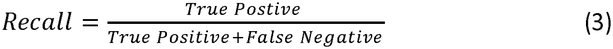

Equation 4: Precision

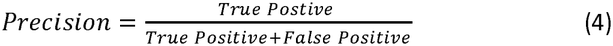

Equation 5: F1 Score

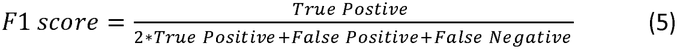

To ensure manual counts by the investigator were accurate and reproducible, a second investigator counted SGNs for six cases, one randomly selected from each etiologic category, using Fiji (ImageJ) (34). Manual counts for these cases were compared between investigators.

## Results

### Developing the Deep-Learning Model

Due to the diverse appearances and densities of SGNs resulting from factors including storage times, staining protocols, post-mortem autolysis, and neuronal degeneration, three segmentation methods were tested prior to creating ground truth annotations to train the model (Fig. 3a). Consistent with established criteria when using eSlides, an SGN was defined by the presence of a visible nucleus within the cytoplasm (25). As done with previous studies, an initial approach included segmenting only the nuclei (25,27). However, this method resulted in overcounting by misclassifying Schwann cells, vessels, and artifacts as SGN nuclei (Fig 3b.1). A second approach included cytoplasmic segmentation, which led to undercounting, as adjacent SGNs were fused into a single detection (Fig. 3b.2). For the third approach, both the nucleus and cytoplasm were segmented. Incorporating both features provided more contextual information for the model, improving feature discrimination and yielding improved and consistent results across cases (Fig 3b.3). From these findings, three classes, nucleus, cytoplasm, and background were segmented to create ground truth data to train the model.

**Fig. 3.**
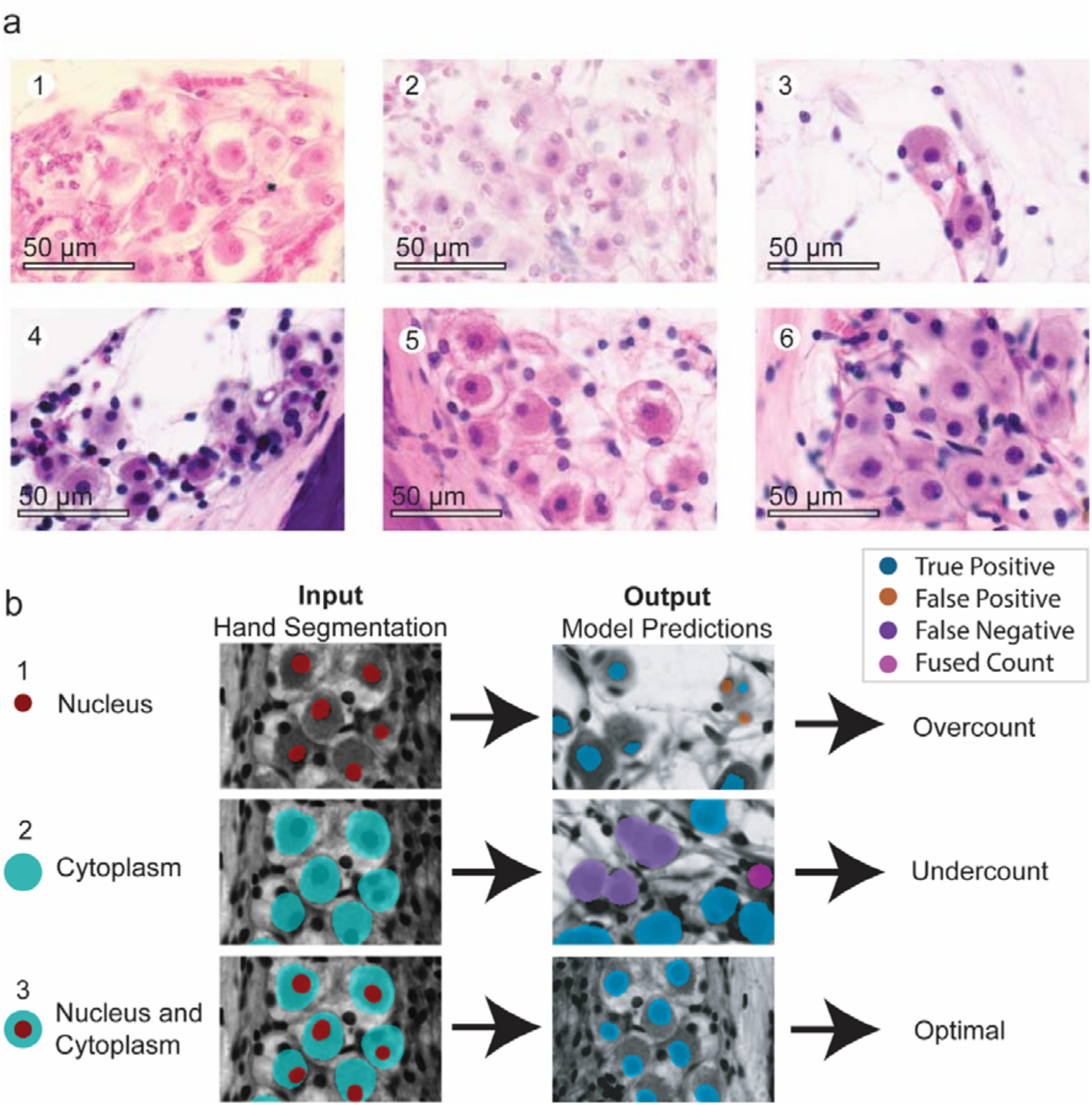
Segmentation methods. Magnified images of SGNs from specimens used to train the model, demonstrating the wide variety of SGN appearances in the lower basal turn (a). A prolonged post-mortem time in which the nuclei appear pycnotic (a.1); relatively faint staining intensity (a.2); significant neuronal degeneration (a.3); cysts within Rosenthal’s canal, causing clumping of SGNs (a.4); a prolonged post-mortem time in which the cytoplasm is condensed (a.5); a short post-mortem time and higher neuronal density (a.6). Comparison of different segmentation methods used to train the model (b). Segmenting the nuclei, led to overcounting (false positives in orange); segmenting the cytoplasm led to undercounting (false negatives in pink and fused counts in purple); segmenting both the nucleus and cytoplasm led to the most true positives (blue).

The U-Net Model was trained using six hTB specimens and both training loss and validation loss decreased over training epochs. Following an initial rapid decrease in the loss values, they plateaued, demonstrating the model learned how to distinguish fundamental morphological features of SGNs.

The Dice coefficient was used to evaluate the overlap between model predictions and ground truth segmentations, which is suitable for multiclass segmentation with imbalanced distributions. The model achieved a Dice coefficient of 0.91 on the validation set.

### Model Accuracy Across Cases

Eighteen independent cases were used to assess the accuracy of the model, consisting of three cases from each of the six diagnoses used to train the model with varying post-mortem and storage times. For each case, three slides were selected from standardized regions throughout the cochlea, representing seven different anatomical areas ranging from the hook region in the base to the apex. These areas enabled assessment of the model’s robustness throughout the length of the cochlea and among different neuronal densities. Precision and recall across all cases were 0.96 and 0.87, respectively, with an F1 score of 0.91 (Fig. 4 and Supplemental Fig. 1).

**Fig. 4.**
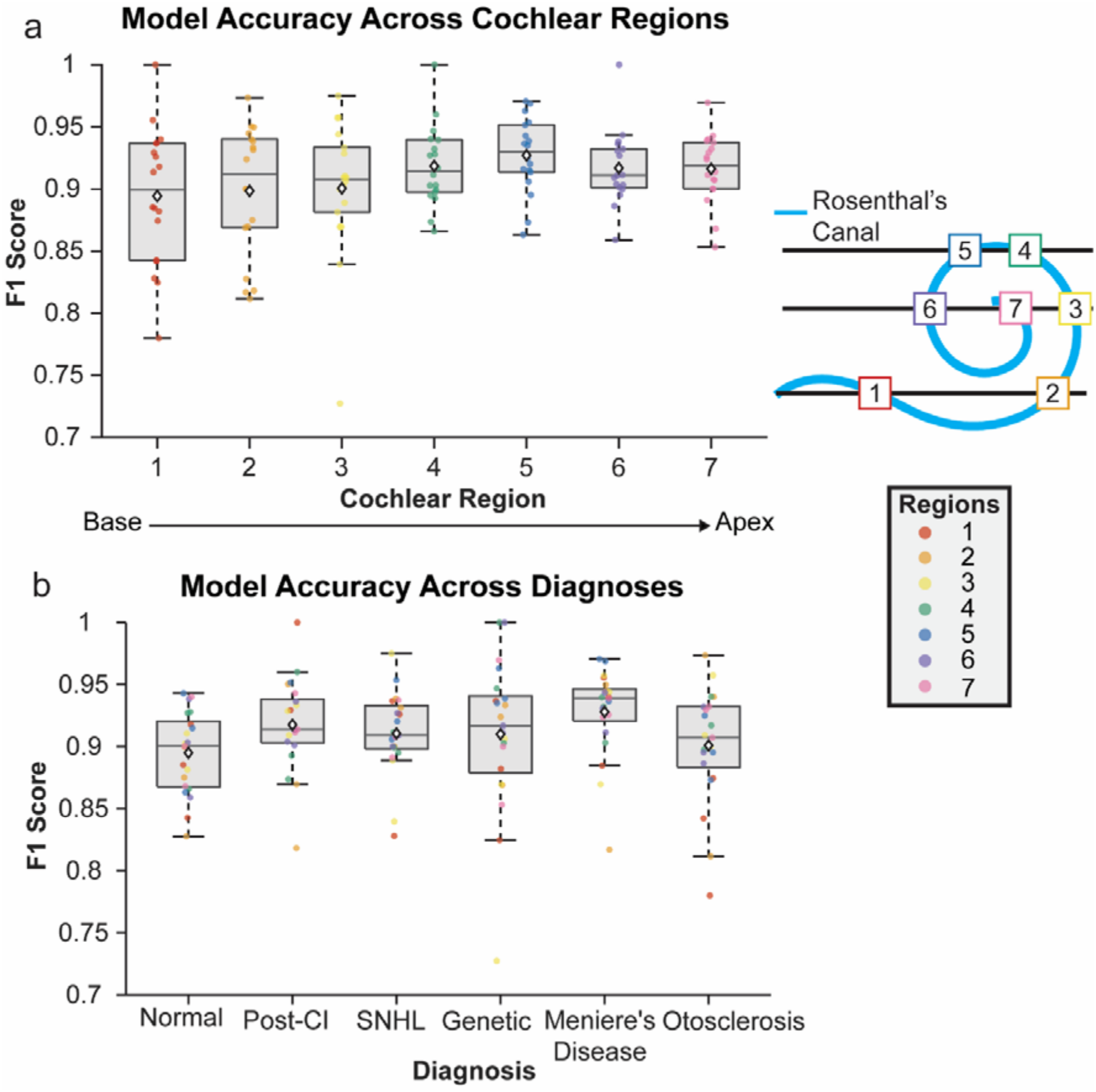
Model performance across eighteen diverse human temporal bone specimens. Model performance across seven standardized regions spanning from base to apex, selected using three histological slides per specimen. Each colored data point represents a corresponding cochlear region (a). Model performance across six different diagnostic categories, including three specimens per etiology (b).

To ensure the manual counts, considered the “gold-standard,” were accurate, a second investigator was blinded to the prior quantification data and re-counted 6/18 cases (one for each hearing loss etiology). The intraclass correlation coefficient (ICC) (A.1) was 0.987 (95% confidence interval: 0.977-0.993, p<0.0001). Bland-Altman analysis showed no systemic bias (intercept p=0.64) and no proportional bias (slope p=0.13). The counts by the investigators showed excellent agreement and reproducibility between the observers was high. Therefore, the remaining 12/18 cases were not manually counted a second time.

### Model Accuracy Throughout the Cochlea

For one hTB specimen, all available slides through Rosenthal’s canal that were stained using H&E were analyzed to further evaluate model performance throughout the entire cochlea (Fig. 5a). Each SGN was labeled as a true positive, false positive, or a false negative and was analyzed from base to apex by percent distance along Rosenthal’s canal. Model counts were highly correlated with manual counts (r^2^=0.99) and the model identified SGNs were accurate in comparison to manually identified SGNs with and F1-score of 0.96 (Fig. 5b). There were a higher number of false positives and false negatives near the hook region and in areas where there were increased neuronal density, such as near the apex (Fig. 5c,d). Using coordinates output by the model, SGNs were plotted and frequency mapped, generating a three-dimensional reconstruction (Fig. 6).

**Fig. 5.**
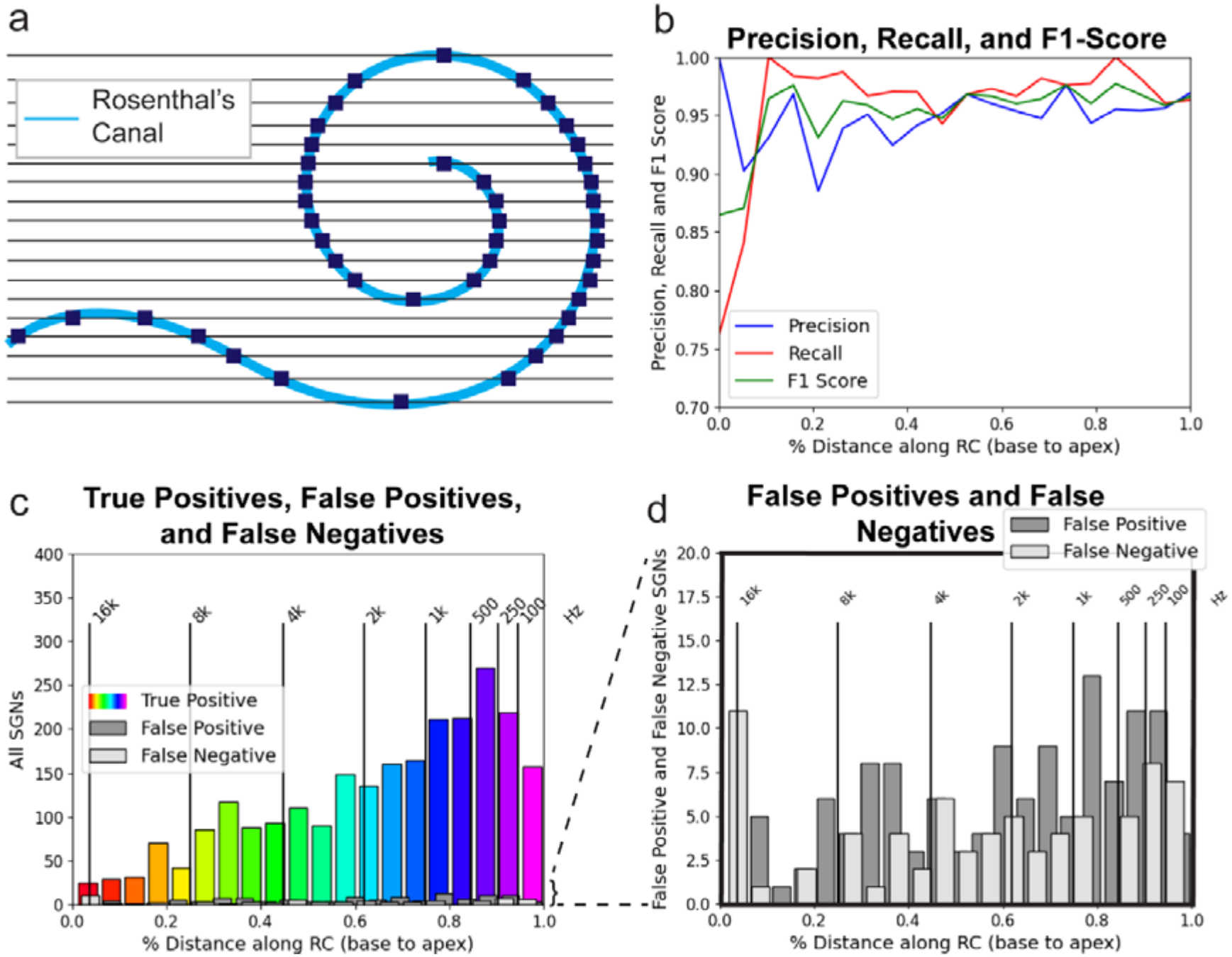
Comprehensive analysis of model performance throughout the cochlea in a single specimen. All available H&E-stained slides including Rosenthal’s canal used to assess the model with the navy boxes indicating areas where performance was assessed (a). Precision, recall, and F1 score plotted from base to apex (b). The number of true positives (color-mapped by frequency), false positives (gray), and false negatives (white) plotted by percent distance along Rosenthal’s canal (c). A magnified view of panel (c) highlighting the number of false positives and false negatives (d).

**Fig. 6.**
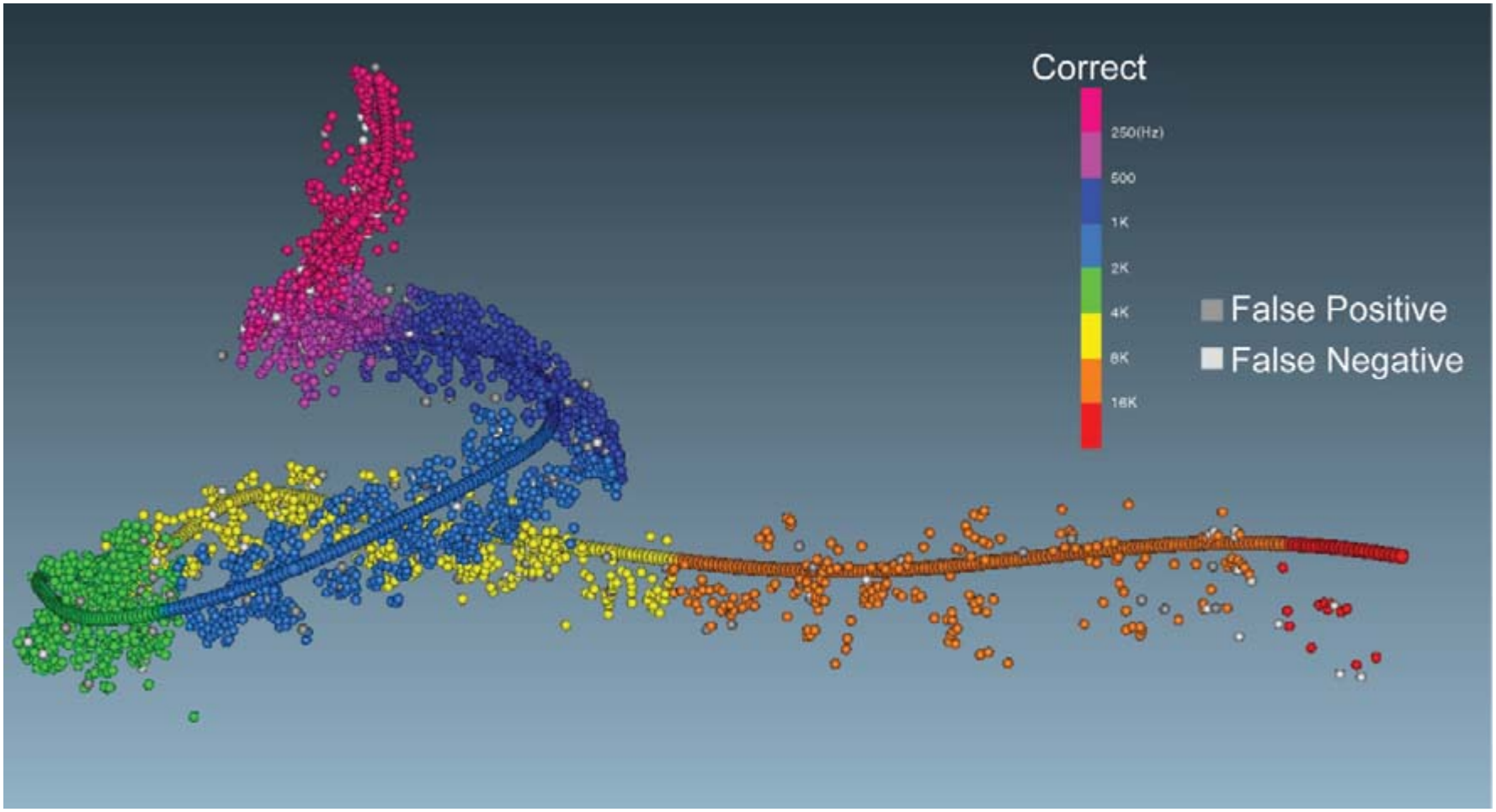
Mapping SGNs. A three-dimensional reconstruction of Rosenthal’s canal with each SGN labeled as true positive, false positive, or false negative plotted from base to apex.

### Model Adaptability

To test the adaptability of the model, the case with the lowest performance, Case A, was used to generate additional training data. A few small areas selected from this case, ranging from base to apex, were hand-segmented, as previously described, and were used as additional input. Fine-tuning took approximately 20 minutes, and this new fine-tuned version of the model was applied to Case A. Model predictions were compared to manual counts. F1 score improved from 0.88 to 0.94, and the r^2^ value improved from 0.919 to 0.997 (Fig. 7a).

**Fig. 7.**
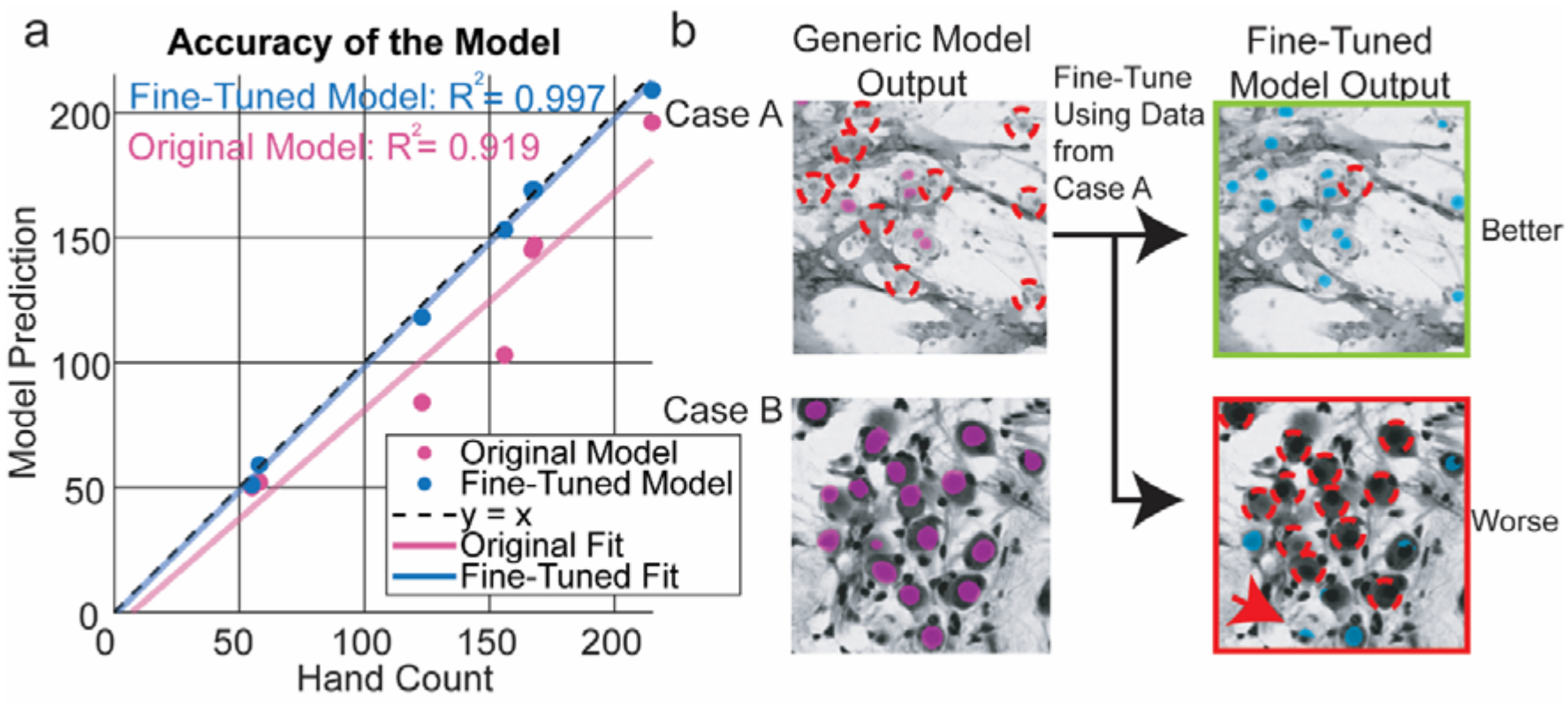
Fine-tuning the model. Accuracy of the original model (pink) and a fine-tuned version of the model (blue) trained by using training data from case A (a). Histological images showing the output of the original model (pink) and the fine-tuned model (blue) for case A (used to train the fine-tuned version) and case B (a new case).

To assess whether overall model performance improved for all cases with additional training data, the fine-tuned model was reapplied to Case B, for which the original model had performed well (Fig. 7b). In case B, SGNs exhibited a well-defined nuclei and cytoplasm, whereas in Case A, the specimen used for fine-tuning, the SGNs were fainter and less distinct. When applied to case B, the fine-tuned model performed poorer than the original model, correctly identifying fewer SGNs.

## Discussion

### Model Performance

We present a novel approach for quantifying SGNs in human cochleae by applying an adaptable deep learning model to histologically processed hTBs. This approach improves efficiency while maintaining high accuracy in comparison to manual quantification by experienced investigators.

The SGN Base Model is widely applicable to heterogeneous cases and cochlear regions. The model accurately identified SGNs across variable hTB specimens with diverse etiologies, cellular morphologies, and neuronal densities, achieving an F1 score of 0.91. Additionally, when applied to every available slide throughout the cochlea for a single hTB case. The model achieved an F1-score of 0.96, demonstrating strong performance across cochlear turns. These results demonstrate the model’s overall robustness across diverse cochlear regions, neuronal densities, and cellular morphologies.

Previously published U-Net models used for histological datasets typically achieve F1-scores of approximately 0.80-0.87 with optimized architectures achieving up to 0.90-0.93 (25,27,35–38). When compared to these previously published deep-learning models, the SGN Base Model performed comparably and achieved accuracy within previously published ranges, indicating that it meets current methodological standards, despite morphological and staining heterogeneity.

The SGN Base Model demonstrated higher accuracy in cases with high-contrast staining, well-preserved SGN morphology, and minimal post-mortem cellular changes. In areas with substantial background noise, including areas within bone and anatomically complex regions, such as the hook, performance decreased. Accuracy was also affected in areas of high neuronal density, particularly where SGNs were partially concealed by axons, Schwann cells, blood vessels, or histologic artifact. Notably, these limitations are also relevant when manually counting SGNs using digitized slides. Although we refer to manual counting as the “ground truth,” manual quantification is susceptible to human error, including missed SGNs that are embedded within the bony regions of the modiolus. Furthermore, it can be difficult to distinguish SGNs in denser areas, especially using digitized slides where there is a single focal plane, which precludes assessment across the full z-depth axis.

### Hand Correction

When extracting the nuclei as a distinct ROI, the Connect Components algorithm in Dragonfly 3D World calculated surface area, which was ultimately used to determine a size cut off. Using a histogram to analyze surface area, a size cut off was generated to eliminate background noise, especially extremely small, potentially mislabeled pixels. In this dataset, and based on a previous study, 0.004 mm^2^ was determined to be a suitable threshold; however, depending on different factors, such as staining techniques, specimen processing, and resolution, a different size cut off may be beneficial (27).

Although the SGN Base Model demonstrates strong performance, we recommend assessing the model’s overall performance after application to a hTB specimen and manually correcting individual slides as needed. Regions of increased background signal and neuronal density demonstrated an increased number of both false positives and false negatives. Additionally, special consideration may be needed to use this deep learning mode for hTB specimens with cochlear implants, as they are associated with intracochlear trauma, such as fibrosis and ossification which can cause difficulty visualizing, and ultimately quantifying SGNs. These considerations may be beneficial when determining areas to spot check for accuracy and evaluating if fine-tuning the model is necessary for the case being analyzed.

### Fine-Tuning and Adapting the Model

The model’s ability to be fine-tuned and improved highlights is adaptability to diverse datasets. Fine-tuning is recommended when accuracy falls below ~80% or when visual inspection of the model’s outputs reveals reduced performance, including specimens with unique staining characteristics and increased background noise.

When fine-tuning using a single hTB case, it is important to include areas representing various cochlear turns and neuronal densities. Furthermore, when creating training data, including background that the model could potentially mislabel as an SGN, such as bone, vessels, and Schwann cells, will improve the model’s ability to discriminate between SGNs and background noise. Because fine-tuning using a single case causes the model to become specialized to the characteristics of that specimen, the fine-tuned version of the model may not be generalizable to all datasets. Therefore, it is recommended to start by applying the original version of the model to each new case. The exception is when a new case exhibits staining intensity or SGN morphology similar to that of a case previously used for fine tuning; in these instances, the fine-tuned version may outperform the original SGN Base Model.

The SGN Base Model is accurate and widely applicable to the eSlides in the Mass Eye and Ear Otopathology Laboratory Collection. While the specimens in other collections may differ due to processing and staining techniques, in a separate study, the model was successfully applied to the digitized slides of two other collections to quantify SGNs in a multi-institutional study. If fine-tuning is necessary to adapt the model to a new dataset, including multiple hTBs with variable post-mortem times, storage duration, neuronal densities, cellular morphologies, and staining intensities may improve the model’s performance across the entire collection.

## Conclusion

SGN quantification in hTBs has been historically a very time and labor-intensive methodology. The SGN Base Model substantially improves efficiency while maintaining high accuracy comparable to techniques considered to be the “gold-standard.” Its adaptability through fine-tuning allows for iterative improvement and broad applicability to diverse datasets. Furthermore, the model’s ability to produce spatial coordinates for each identified SGN enables detailed mapping of neuronal density from base to apex. As such, this approach could be used to investigate hearing loss of diverse etiologies and has the potential to inform the development and optimization of therapeutic strategies in which it is essential to understand the distribution of SGNs throughout the cochlea, such as cochlear implantation.

## Acknowledgements

The authors are grateful to all the donors and their families for donating and assisting with the donations of temporal bones through the NIDCD National Temporal Bone Registry. We thank MengYu Zhu, Diane Jones, and Barbara Burgess for processing the histological specimens used in this study and Paula Robles for discussion about statistics.

## Statements

### Statement of Ethics

#### Study approval statement

This study was reviewed and approved by the Institutional Review Board of Massachusetts Eye and Ear (IRB #2021P001593).

#### Consent to participate statement

Consent for the temporal bone specimens included in this study was given by the donors or their legal representatives.

### Author Contributions

AKH, AMQ, and JGA designed this study. AKH, HW, and CG contributed to model development. AH and NK contributed to analyses. AMQ and JGA oversaw the project. All authors reviewed and approved the final manuscript.

### Data Availability Statement

The code used and developed for this study are available on GitHub.com. Data and support to assist with using the SGN Base Model are available upon request.

## Figure Legends

**Supplemental Fig. 1.**
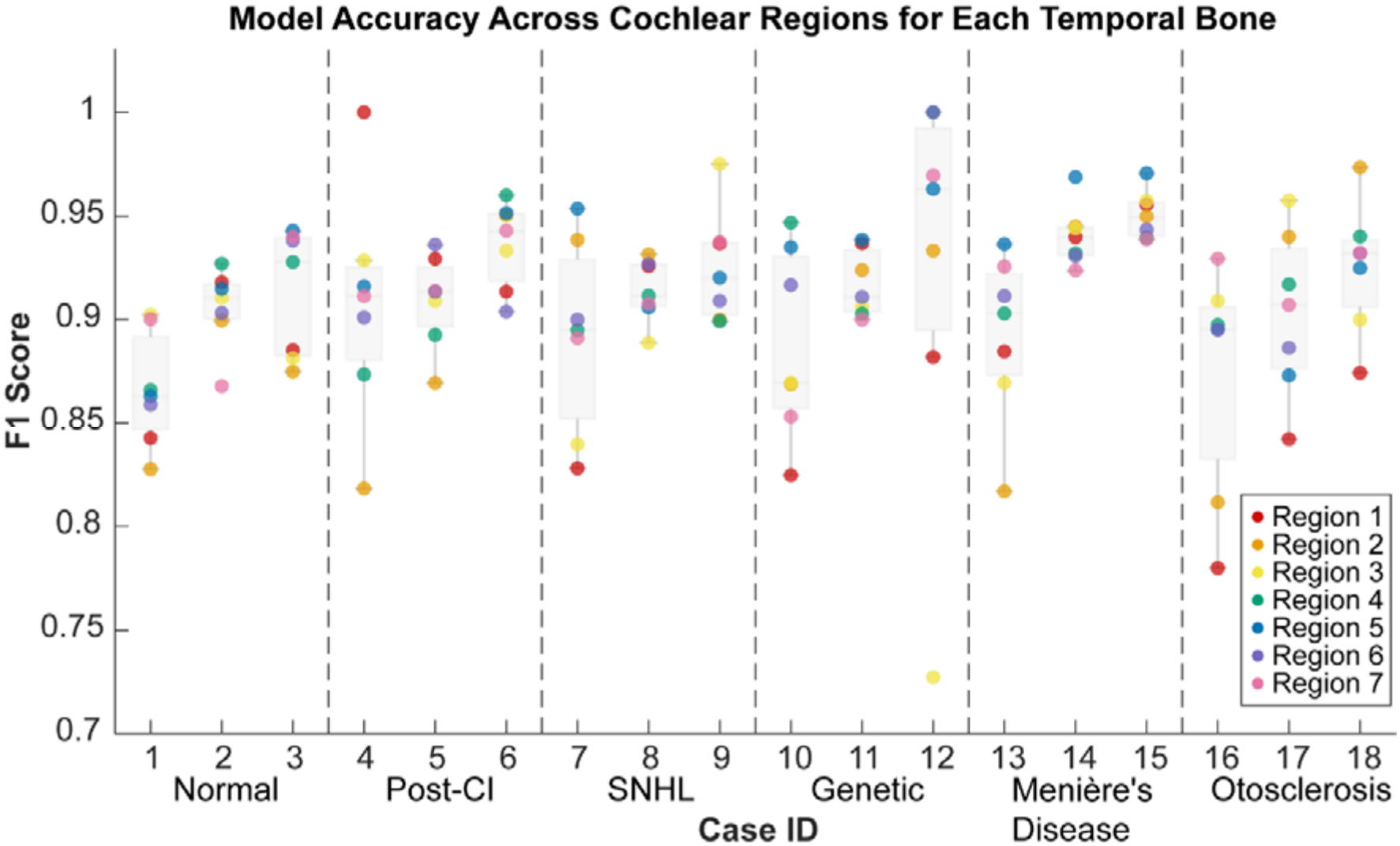
Model performance across the seven standardized regions in eighteen human temporal bone specimens. Each colored point represents the F1 score for a specific predefined region within a given case. Cases are grouped by diagnosis, with dashed vertical lines indicating transitions between diagnostic categories.

